# A dedicated caller for *DUX4* rearrangements from whole-genome sequencing data

**DOI:** 10.1101/2024.05.23.595509

**Authors:** Pascal Grobecker, Stefano Berri, John F. Peden, Kai-Jie Chow, Claire Fielding, Ivana Armogida, Helen Northen, David J. McBride, Peter J. Campbell, Jennifer Becq, Sarra L. Ryan, David R. Bentley, Christine J. Harrison, Anthony V. Moorman, Mark T. Ross, Martina Mijuskovic

## Abstract

Rearrangements involving the *DUX4* gene (*DUX4-*r) define a subtype of paediatric and adult acute lymphoblastic leukaemia (ALL) with a favourable outcome. Currently, there is no ‘standard of care’ diagnostic method for their confident identification. Here, we present an open-source software tool designed to detect *DUX4*-r from short-read, whole-genome sequencing (WGS) data. Evaluation on a cohort of 210 paediatric ALL cases showed that our method detects all known, as well as previously unidentified, cases of *IGH::DUX4* and rearrangements with other partner genes. These findings demonstrate the possibility of robustly detecting *DUX4*-r using WGS in the routine clinical setting.

## Background

In childhood and adult B-cell acute lymphoblastic leukaemia (B-ALL), chromosomal abnormalities play a significant role in risk stratification for treatment within clinical trials worldwide. Although a wide range of genetic aberrations have been known for many years, recent sequencing studies have uncovered a wealth of additional genetic information of prognostic relevance with implications for changes to treatment strategies. One such genetic alteration, *DUX4* rearrangements (*DUX4*-r), defines a recently reported subtype that affects 4–7% of paediatric patients [1] and ∼5% of adolescents and young adults [2]. Patients in all age groups exhibit favourable outcomes but due to the presence of concomitant risk factors are frequently treated as intermediate or high risk [3,4]. However, for reasons described below, the detection of *DUX4-r* is challenging and if optimal therapy is to be given to these patients, their accurate identification is paramount.

DUX4 (Double Homeobox 4) is a transcription factor that is selectively and transiently expressed in cleavage-stage embryos [5] and germ cells of the testis [1]. A copy of the *DUX4* gene, encoding two homeoboxes, is located within each unit of the D4Z4 macrosatellite repeat array in the subtelomeric region of chromosome 4 long arm (4q) and in a similar repeat array on chromosome 10q. The ∼3.3 kb D4Z4 repeat is polymorphic in length and has 11-100 copies in healthy individuals [6]. When ectopically activated, DUX4 can upregulate expression of multiple genes and initiate transcription from alternative promoters, leading to non-canonical transcript isoforms [7]. In fact, it has been shown that contraction of the D4Z4 repeat array below 11 copies decreases the epigenetic repression of *DUX4*, causing autosomal dominant facioscapulohumeral muscular dystrophy (FSHD) [8].

*DUX4*-r cases in ALL were initially discovered through their distinctive gene expression profile [9]. *DUX4* is commonly rearranged with the Immunoglobulin Heavy Chain Locus (*IGH*), although multiple fusion partners have been identified [10–14]. The rearrangement typically creates a chimeric transcript that retains the 5’ end of *DUX4* but replaces the 3’ coding sequence with a section of *IGH*. This event, most likely via *IGH* enhancer hijacking, leads to activation of expression of *DUX4* in developing lymphocytes. The resulting change in the transcriptional landscape of the affected cell is thought to lead to oncogenic transformation [13]. *DUX4*-r have also been discovered in a distinct rare subtype of *CIC* (Capicua Transcriptional Repressor)-rearranged non-Ewing sarcoma, accounting for less than 1% of all sarcomas [15], primarily affecting young adults and associated with a poor outcome [16]. In *CIC::DUX4* sarcoma, similar to ALL, the resulting fusion protein acts as a transcriptional activator driving the oncogenesis [17].

In ALL, *DUX4*-r were first thought to be driven by co-occurring deletions in the ERG transcription factor, which was used as a surrogate for their identification. However, more recent studies have now indicated that *ERG* deletions are present in only a subset of *DUX4*-r patients and that they are likely to be subclonal [3,12,14,18,19]. Subsequently, using next generation sequencing (NGS) approaches, two independent studies confirmed *DUX4*-r to be the driving lesion [11,13].

The complex and cryptic nature of *DUX4*, and the presence of very similar *DUX4* copies throughout the genome, have precluded its accurate detection using current bioinformatics tools and other standard of care genetic tests. Common approaches that look for accumulation of discordant sequence read-pairs to identify breakpoints typically fail due to the multiple possible mapping locations leading to scattering of supporting reads. Furthermore, most structural variant callers disregard sequence reads with multiple equivalent mapping positions. Clearly, robust genetic testing methods are urgently needed. We reasoned that a custom *DUX4*-r caller, which takes into account all read-pairs spanning any *DUX4* copy and the gene partner of interest, was required: an approach that we have described previously [12]. We present here an improved implementation of this method as Pelops, an open-source software tool that can be integrated into existing bioinformatics analysis pipelines. We evaluated Pelops on a paediatric B-ALL cohort of 210 patients [12] and demonstrate that Pelops is a robust tool for identifying *DUX4* rearrangements from tumour-only WGS data. This proof-of-concept work indicates a path to improved diagnostic testing for this good risk genetic subtype in clinical WGS pipelines.

## Results

### Pelops overview

Pelops is a software tool that implements a tumour-only analysis to identify the signal of *DUX4* rearrangements from short-read WGS data, building on our earlier version of the method [12]. The most common *DUX4*-r*, IGH*::*DUX4,* is identified using a targeted approach by finding read-pairs spanning any part of the *IGH* and *DUX4* regions and then calculating the number of spanning read pairs per billion total reads (SRPB) (Figure 1A, Methods, Supplementary Figure S1).

**Figure 1.**
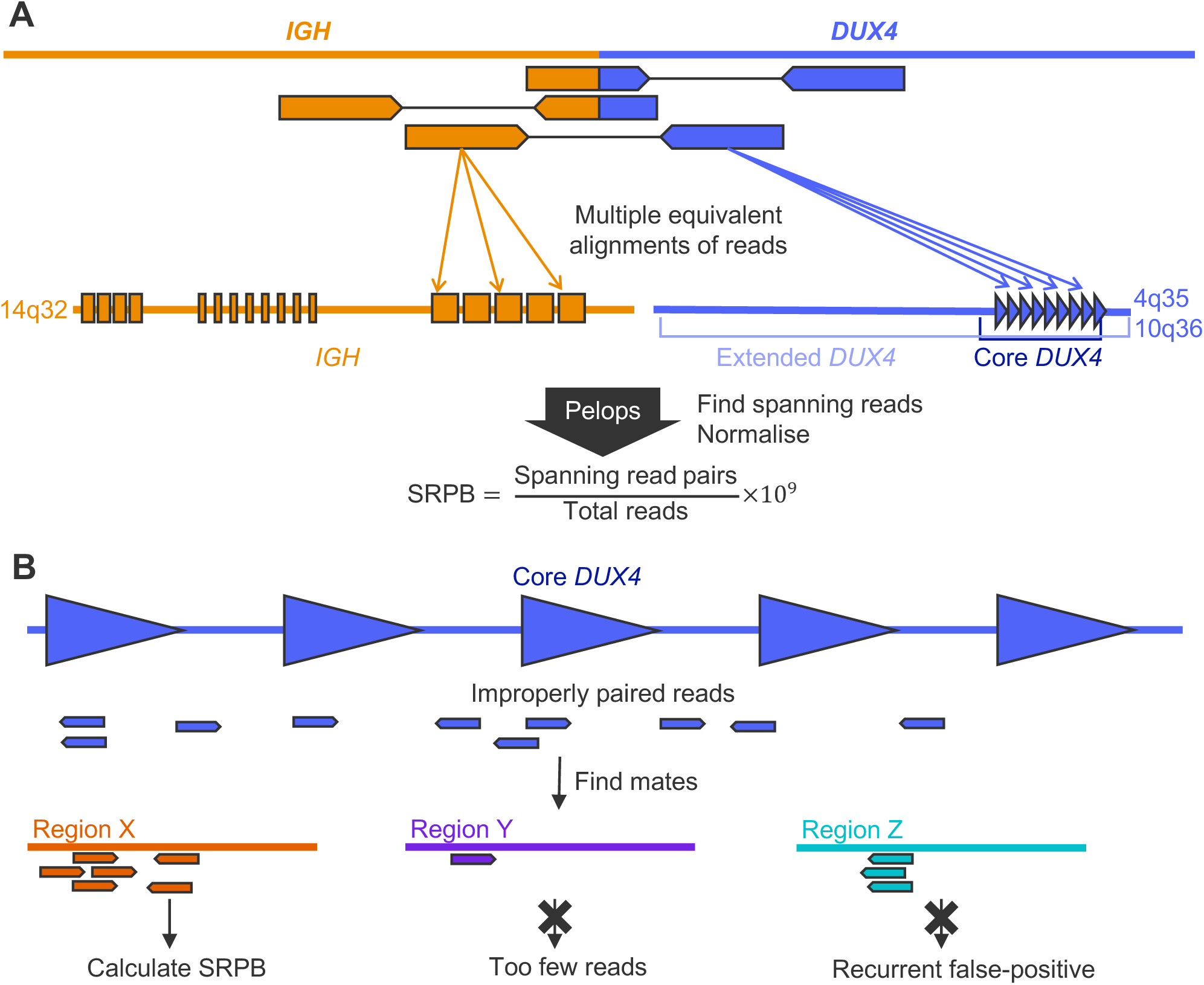
Overview of Pelops’ *DUX4*-r detection method. (A) Detection of *IGH::DUX4* fusions. Reads spanning the *IGH::DUX4* translocation breakpoint contain segments that align to *IGH* and *DUX4*. These segments are frequently aligned to several different repeats. Pelops finds spanning reads across the whole *IGH* and *DUX4* regions and normalises their count to spanning read pairs per billion (SRPB). (B) Detection of other *DUX4* rearrangements. First, Pelops identifies improperly paired reads in the core *DUX4* region, with mates mapping anywhere else across the genome. Then, regions where multiple mates are clustering are identified. Finally, any regions with the number of mates below a threshold, as well as recurrent false positive regions are removed. SRPB values are calculated for all remaining regions.

For *DUX4*-r cases involving *DUX4* fusions with other partner genes, an untargeted, genome-wide approach is required. Pelops finds evidence for these cases by identifying genomic regions containing multiple mates of improperly paired reads anchored in the *DUX4* region (Figure 1B, Methods, Supplementary Figure S1).

### Evaluation on the paediatric ALL cohort

#### *IGH::DUX4* rearrangements

An earlier implementation of this approach identified 57 *IGH::DUX4* cases in a cohort of 210 paediatric B-cell ALL patients [12] (Supplementary Table 1). As described in the Methods section, we modified the original method to find all spanning reads in each sample and developed the Pelops software. Initially, we ran Pelops on the same 210 leukaemia samples, using their matched germline samples as negative controls. We found excellent concordance between published spanning reads per billion (SRPB) from the earlier implementation with those from Pelops (Supplementary Figure S2). The SRPB values depend on how the *DUX4* region is defined, and we show results for two complementary definitions: the ‘core *DUX4*’ region and the ‘extended *DUX4*’ region. The extended region covers the *DUX4* repeat arrays on chromosomes 4 and 10 with a 100 kb margin, as well as additional *DUX4* pseudogenes on other chromosomes. The core region is a subset of the extended region, and only covers the subtelomeric *DUX4* repeats on chromosomes 4 and 10, with a 1 kb margin (Figure 1A, Supplementary Table 2). By using SRPB values calculated for these two regions, it was possible to identify all samples with an *IGH::DUX4* fusion (Figure 2). These results were independent of the read aligner used: while we discuss results for DRAGEN alignments (Figure 2) for the remainder of this section, bwa and Isaac [20,21] yielded qualitatively identical results (Supplementary Figure S3, Supplementary Table 3).

**Figure 2.**
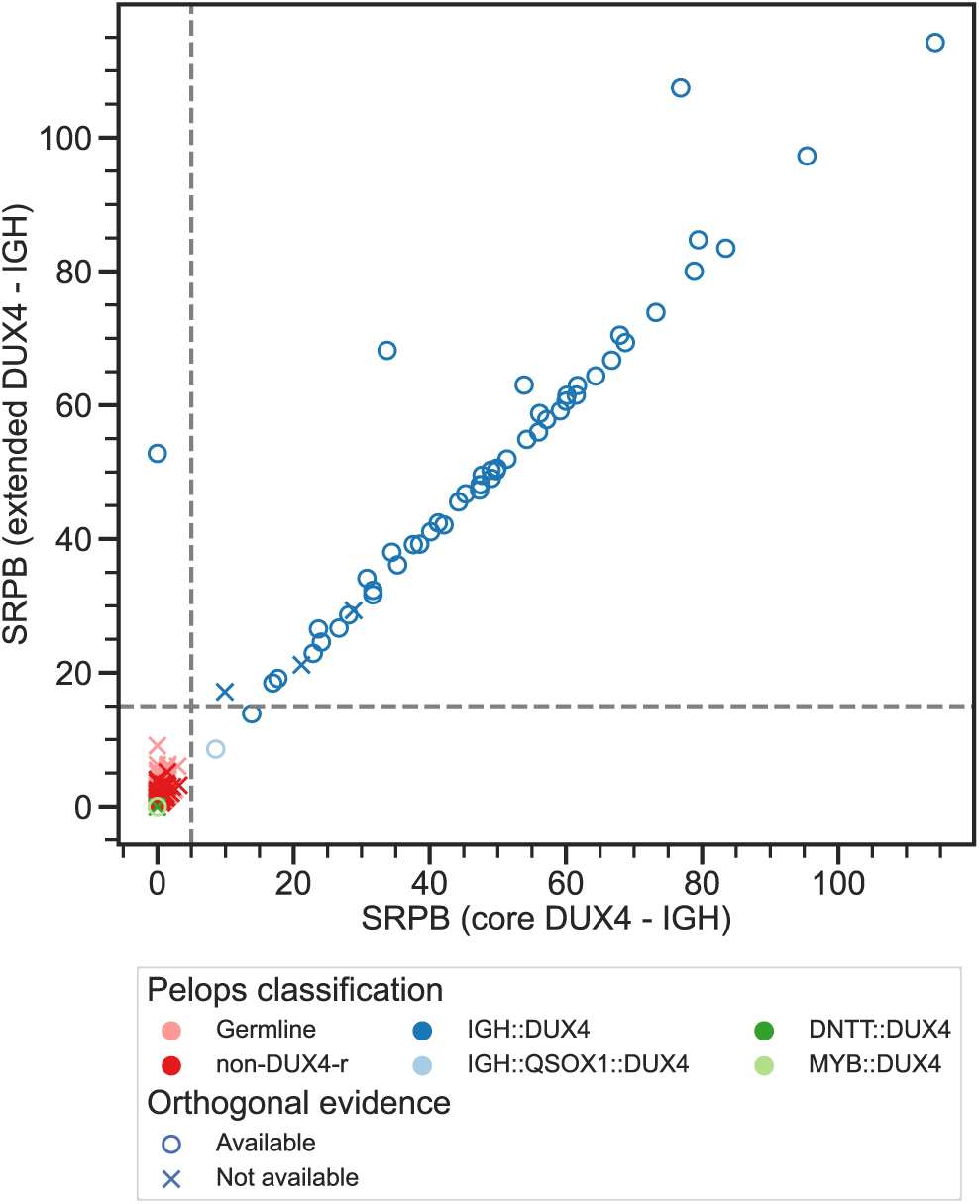
Spanning read pairs per billion (SRPB) for *IGH::DUX4* calculated by Pelops for all 210 tumour and 208 matched germline samples of the evaluation cohort, using alignments by DRAGEN. The x-axis shows the SRPB distribution calculated based on the core *DUX4* region definition, while on the y-axis calculations are based on the extended *DUX4* region definition. Colours indicate *DUX4* fusion type of the samples predicted by Pelops. The dashed horizontal and vertical lines indicate SRPB thresholds of 15 and 5, for extended and core *DUX4* regions, respectively, on which the identification of *IGH::DUX4* fusions is based. The marker indicates whether orthogonal evidence for *DUX4*-r is available, based either on RNA-sequencing (gene expression profile and/or *IGH*::*DUX4* fusion), the presence of *ERG* deletions, or amplicon sequencing.

Among the 57 samples identified by Pelops as *IGH::DUX4* fusions, orthogonal evidence was available for a total of 55 (Figure 3, Supplementary Table 1). Previous analysis confirmed 43 samples through *ERG* deletions, and through whole-transcriptome sequencing (WTS) [12]. *ERG* deletions associated with *DUX4*-r were confirmed by WGS and by Multiplex Ligation-dependent Probe Amplification (MLPA). Whole-transcriptome sequencing (WTS) was used to confirm cases based on their gene expression profile and presence of *IGH::DUX4* RNA fusions. In this study, we used Pelops outputs to create *de novo* sequence assemblies of *DUX4*-r breakpoint junctions, as described in more detail below. Based on these sequences, we then designed primers to confirm a further 12 *IGH::DUX4* cases by PCR amplicon sequencing (Figure 3, Supplementary Table 1).

**Figure 3.**
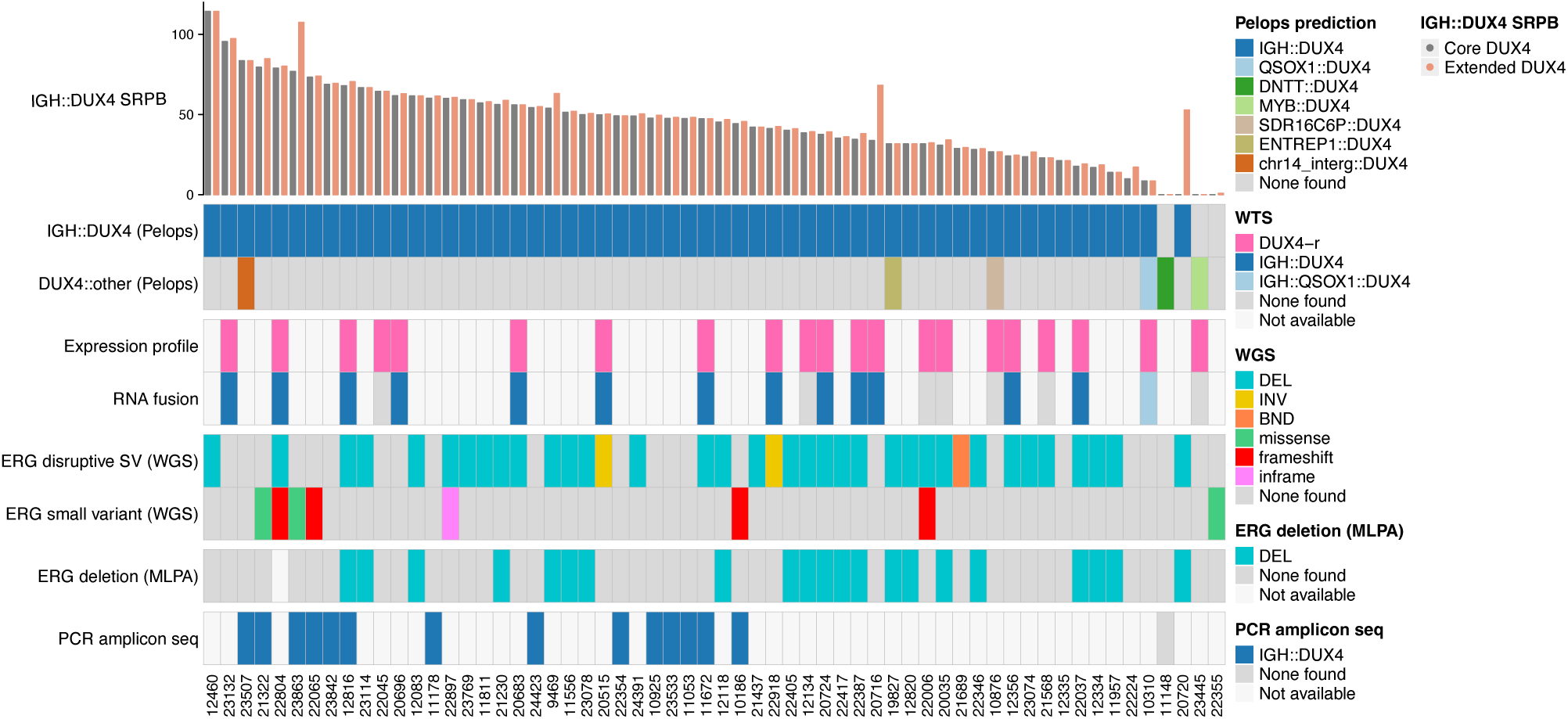
Summary of orthogonal evidence for *DUX4* rearrangements available for samples classified as *DUX4*-r in Ryan et al, 2023 and evidence from Pelops. Patient IDs are shown at the bottom of the plot.

Using the core *DUX4* region, all but one of the orthogonally validated samples were correctly called using a threshold of SRPB ≥ 5 (Figure 2). The remaining case (patient 20720) was called using the extended region as described below. Of the 21 *IGH::DUX4* samples with relatively low SRPB (5 ≤ SRPB < 40), 18 had orthogonal evidence of *DUX4*-r (Figure 3). The level of noise in germline and leukaemia samples with no known *IGH::DUX4* fusion was very low, with SRPB < 3.1 in the core region. Thus, we are confident that Pelops has a precision of 100% in this cohort, at a threshold of SRPB ≥ 5 for DRAGEN alignments in the core *DUX4* region.

The paediatric validation cohort contained only one case (patient 20720) with an *ERG* deletion but no spanning reads between the core *DUX4* region and *IGH* (Figure 2). The *DUX4*-r was successfully identified (SRPB = 52.8) only by using the extended *DUX4* region that includes sequence 100 kb upstream of *DUX4*. In other samples, using the extended *DUX4* region also uncovered additional breakpoints (Supplementary Figure S6). However, it also led to an increase in false positive spanning reads in germline samples (e.g. patient 21322; SRPB = 9.1). This necessitated an increase in the threshold to SRPB ≥ 15 when using the extended region definition.

The number of spanning reads defining a genuine *IGH::DUX4* rearrangement should theoretically depend on several sequencing and sample parameters. A longer read length should increase the split read evidence, while a larger insert size should increase the paired read evidence. Higher tumour content and sequencing coverage should lead to an increase in both types of spanning reads. There is a clear correlation between all these parameters with SRPB (Supplementary Figure S4). While the SRPB measure is normalised for the number of reads, sequencing parameters changed over the course of the paediatric ALL study and tumour samples sequenced earlier had lower coverage, read length and fragment size compared to those sequenced later (Supplementary Figure S5). As a result of this interdependency, it was not possible to separate the effects of each of the parameters on SRPB. However, the data in Supplementary Figure S4 show that Pelops called *IGH::DUX4* rearrangements in the most challenging samples of the cohort, which have 100 base read-length (vs 150 base in most samples), short median insert size (< 300 bp), low coverage (30-40×), or low tumour content (< 50%). Finally, the choice of aligner also has a clear impact on the SRPB values as they are sensitive to the aligner’s behaviour in challenging-to-map regions (Supplementary Figure S3, Supplementary Table 3). For reads aligned with bwa, SRPB values are higher on average than with DRAGEN, while Isaac alignments yield lower values. Considering these factors, it is remarkable that simple fixed thresholds of SRPB ≥ 5 for the core *DUX4* region and SRPB ≥ 15 for the extended *DUX4* region consistently yield correct results in this cohort irrespective of the aligner used.

#### *DUX4* rearrangements with other partner genes

The earlier analysis of the paediatric B-ALL cohort [12] identified two *DUX4-*r in which *DUX4* expression was possibly activated through a fusion with non-*IGH* genes active in developing lymphocytes, *MYB* and *DNTT*. We further improved and automated the method for calling such rearrangements (Figure 1B), as described in more detail in the Methods. Pelops was run on all leukaemia samples from the paediatric cohort and was able to recall the *MYB::DUX4* and *DNTT::DUX4* rearrangements (Figure 3). It also called a *QSOX1::DUX4* rearrangement which is part of an *IGH::QSOX1::DUX4* triple fusion. Three more, previously undescribed, rearrangements were called in samples which also contained an *IGH::DUX4* fusion. In patients 10876 and 19827, the translocations were found in introns of *SDR16C6P* and *ENTREP1*, respectively. In patient 23507, the rearrangement did not fall inside a gene or pseudogene. Furthermore, despite the presence of a somatic *ERG* missense variant, we found no evidence for a *DUX4*-r in one case (patient 22355) using Pelops, which is in agreement with the previous implementation.

### *De novo* assembly and validation of *DUX4*-rearrangement breakpoints

To provide additional confidence in this method, we sought to confirm that the spanning reads identified by Pelops could be used for a *de novo* assembly of the translocation breakpoints. For all 59 *DUX4*-r cases, we were able to obtain at least one sequence assembly that aligned to a relevant breakpoint junction (Methods, Supplementary Table 4). All six cases of *DUX4*-r with non-*IGH* regions were also confirmed in this way. This provides an important additional validation of the predicted *DUX4-*r and confirmed that Pelops made no false positive calls within the paediatric patient cohort.

Amongst the 57 samples with *IGH*::*DUX4* rearrangements, we were able to find two or more breakpoint junctions in 67% (38/57) of cases (Supplementary Figure S6). Due to the repetitive nature of the *DUX4* region, it was not always possible to distinguish between an insertion of a *DUX4* sequence into *IGH*, a reciprocal translocation event, or multiple independent rearrangements of the two alleles. However, for five samples, the *de novo* assembled sequence captures an entire insertion of a *DUX4* sequence into the *IGH* locus, which has been described as the most common mechanism for creating a *DUX4* fusion that acts as an oncogenic transcriptional activator [1].

When looking at the genomic positions of breakpoints, we observed a clear association with V, D and J segments of the *IGH* gene (Supplementary Figure S7), the targets of V(D)J rearrangements during lymphocyte development. Most breakpoints were clustered in a 60 kb genomic region containing IGH-D and IGH-J segments, which makes up less than 5% of the entire *IGH* region.

To obtain orthogonal evidence and confirm Pelops results for the 16 *DUX4*-r cases that were not validated in the previous study [12], the sequence assemblies described above were used to aid PCR primer design for amplicon sequencing. Out of these 16 samples, DNA was available for 13. Two additional samples, previously confirmed through RNA-sequencing, were used as positive controls, bringing the total number of samples used in this experiment to 15. A unique PCR product was obtained in 14 cases (Supplementary Table 1, Figure 3). PCR primer design was not possible on the remaining case due to the repetitive nature of the rearrangement junction. The PCR amplification products were successfully sequenced, confirming that the *de novo* assembly based on spanning reads identified by Pelops accurately represented the rearranged sequence (Methods).

## Discussion

In the NHS England Genomic Medicine Service, all acute leukaemia cases are eligible for WGS for genetic subtyping to assist in treatment decisions. However, there is currently no available bioinformatics tool that can robustly detect the *DUX4*-r cases from WGS data. The results presented here show that Pelops, our dedicated *DUX4-r* caller, can accurately detect *IGH::DUX4* rearrangements that have been previously identified by one or more of the following: gene expression profiling, PCR-based assays (MLPA), co-occurring *ERG* deletions, and the presence of RNA fusions. Furthermore, in agreement with our previously published study [12], Pelops detected cases of either *IGH::DUX4* or *DUX4* rearranged with other gene partners, that were not identified using current standard-of-care methods. Additional validation of these cases using *de novo* assemblies of supporting sequencing reads identified by Pelops followed by PCR amplicon sequencing provides further confidence by confirming these results. Importantly, there were no false positive cases among 208 germline samples and 151 leukaemia samples from other currently known B-ALL subtypes.

To our knowledge, no other direct or indirect method can robustly detect *DUX4*-r cases. Although the correlation of *DUX4*-r with *ERG* abnormalities, primarily deletions, is known, our data confirm the findings of other studies [3,12,14,18,19] by showing that only 34/57 (60%) of *DUX4*-r cases in our paediatric B-ALL cohort had a concurrent *ERG* deletion. We also observed no clear correlation of missense mutations in *ERG* with *DUX4* rearrangements. Furthermore, while WTS may be used to classify some *DUX4*-r cases from their distinct gene expression profile, RNA sequencing of leukaemia samples is not yet widely implemented in the clinical setting.

We have shown that rearrangement supporting reads identified by Pelops can be used to assemble contigs and/or scaffolds spanning the junction between fusion genes, which provides additional confidence in our tool. These assemblies can be used for further validation by PCR-based approaches and examined to elucidate mechanistic origins of these rearrangements. One intriguing possibility that requires further investigation is involvement of the aberrant RAG1/2 recombinase activity in the mutagenesis process, given the fact that many *IGH* rearrangement breakpoints joining *DUX4* are in the proximity of RAG recombination signal sequence flanking V, D and J segments of the *IGH* variable region. Similar off-target V(D)J rearrangements involving known oncogenes and tumour suppressor genes have previously been described in the *ETV6::RUNX1* subtype of ALL [22] and in mouse models of lymphoma [23].

It was reassuring that Pelops successfully detected *IGH::DUX4* rearrangements from samples that were analysed using a variety of sequencing and library preparation methods, and had variable tumour content. While we do observe some positive correlation of SRPB values with insert size, sequencing coverage and tumour content, the method was robust enough to detect all known cases in the cohort we analysed. Subclonal complexity and low tumour content were not observed in our validation cohort, but these could impair detection and should be considered when developing sequencing coverage targets. We demonstrate that Pelops can be successfully used on data generated by different sequencing read aligners, including the most widely used open-source tool, bwa. However, this study showed that each alignment tool leads to a different level of “background noise”, which can be further complicated by using different versions of the reference genome. While detection sensitivity of *IGH::DUX4* rearrangements can be easily adjusted by shifting the SRPB threshold for calling positive cases, detecting rearrangements of *DUX4* with other gene partners in a genome-wide approach may require both calibrating the SRPB threshold, mapping quality scores of read mates, as well as blacklisting genomic regions that lead to recurrent false positive calls.

Besides its application in ALL, where the majority of rearrangements occur between *DUX4* and *IGH*, we envisage other applications in ALL, as well as other cancers, where *DUX4* is rearranged with other gene partners. Notably, a subtype of non-Ewing sarcoma with *CIC*-rearrangements is known to primarily consist of *CIC::DUX4* fusions, which are similarly difficult to detect with common bioinformatics tools [16]. To accommodate this need, we designed Pelops to detect *DUX4* rearrangements in a gene-partner agnostic approach in addition to the *IGH*-targeted approach. In fact, Pelops has been able to detect an orthogonally validated *CIC::DUX4* fusion in one available case of non-Ewing sarcoma (data not shown). Although validation of the gene-agnostic method is not the subject of this study, we expect that availability of Pelops will catalyse this process in appropriate cohorts and implementation of *DUX4*-r detection in a wider range of tumour types. We also envisage that this approach could be successfully adapted and applied to identify rearrangements in other genes within repetitive regions that are challenging to detect by current workflows.

## Conclusions

Pelops is an open-source software tool designed to detect *DUX4* rearrangements in short read whole genome sequencing data. In our cohort of paediatric ALL samples, Pelops reliably detected all known *IGH::DUX4* rearrangements as well as additional cases, including *DUX4* rearrangements with other gene partners. Pelops is easy to integrate into existing bioinformatics pipelines and supports inputs created from the most widely used alignment tools such as bwa and DRAGEN. The work described here demonstrates that there is a path to using WGS to meet the current need to aid in the diagnosis and clinical management of ALL and other cancer types driven by *DUX4*-r.

## Methods

### Sequence alignment

Sequence reads were aligned to human genome reference GRCh38 using three different workflows:

- Isaac Aligner (version SAAC01325.18.01.29). The full workflow was described previously [12].
- DRAGEN (version 4.0.3).
- bwa mem (version 0.7.17). Following alignment, read pairs were annotated (‘fixmate’) and sorted, and duplicates were marked using samtools version 1.15.1.

### Calling *IGH::DUX4* fusions

In order to call an *IGH::DUX4* fusion, Pelops finds read pairs spanning the translocation breakpoint(s), which can either be paired reads or split reads. In paired reads, one of the reads maps to *DUX4* and the other to *IGH*. In split reads, one of the two paired reads has a split alignment, with the primary segment aligned to *IGH* and a supplementary alignment in *DUX4*, or vice versa. As both *IGH* and *DUX4* have numerous homologous repeats, segments of spanning reads are frequently mapped to different repeats and could be found across the entire *IGH* and *DUX4* regions. Therefore, the following strategy was used to find *IGH::DUX4* fusions:

1. Count reads spanning the *DUX4* and *IGH* regions. Reads flagged as duplicates or QC failed are filtered out. There is no filter based on mapping quality as relevant reads are frequently in regions with low mappability due to the repetitive character of *IGH* and *DUX4*.
2. Normalise the counts to account for sample-specific coverage, to obtain spanning read pairs per billion (SRPB): 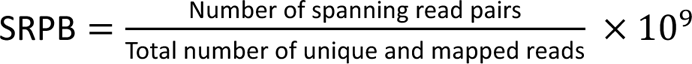.
3. Call *IGH::DUX4* fusion if SRPB is above a given threshold.

Defining the *IGH* and *DUX4* regions. The full definitions of the *IGH* and *DUX4* regions are listed in Supplementary Table 2. The *IGH* region definition was taken from IMGT [24]. To define the *DUX4* regions, several aspects were taken into account:

- *IGH* has a strong enhancer that can activate *DUX4* expression from a long genomic distance, which implies that read evidence for *DUX4*-r could be mapped far upstream of the *DUX4* repeat arrays.
- There are multiple annotated *DUX4*-like pseudogenes outside the *DUX4* repeat arrays. While they are not known to be implicated in *DUX4*-r, spanning reads may still be mapped there.
- A wide region definition could lead to false positive spanning reads due to the presence of repetitive genomic elements.

To accommodate those conflicting requirements, SRPB was calculated based on two *DUX4* region definitions.

The “core *DUX4* region” contains all *DUX4* genes (and pseudogenes) in the repeat arrays on chromosomes 4 and 10 as annotated by Ensembl v91 [25], with a 1 kb margin. It covers a very limited amount of intergenic sequence and leads to few false positive spanning reads in negative samples. Furthermore, this definition is used to call non-*IGH DUX4*-rearrangements with Pelops (see below).

The “extended *DUX4* region” contains the complete subtelomeric repeat arrays on chromosomes 4 and 10 with a 100 kb margin. *DUX4*-like pseudogenes on other chromosomes, as annotated by Ensembl v91, are also included with a 1 kb margin. This region can be used to call rare *IGH::DUX4* rearrangements with breakpoints further than 1 kb upstream of the *DUX4* repeat arrays and rearrangements where the majority of spanning reads were mapped to *DUX4* pseudogenes outside the repeat arrays (although we did not observe such a case in our study). This comes at the expense of having to set a higher SRPB threshold for calling *DUX4*-r, due to the higher rate of false positive spanning reads.

#### Differences to previously published method

While Pelops results are similar to those of the previously published method, three key improvements were made:

1. Counting of all spanning reads. Previously, only reads flagged as improperly paired in the BAM file were used to identify spanning reads. In Pelops, split reads are included which are often flagged as properly paired.
2. Calculation of SRPB. Previously, the numerator of the SRPB equation was based on the number of spanning reads. In Pelops, the calculation is based on spanning read *pairs* to avoid ambiguity in how to count reads with multiple aligned segments.
3. Definitions of *DUX4* and *IGH* regions. Both the *IGH* and the *DUX4* region definitions were changed to reduce noise. The *IGH* region used by Pelops is reduced by 50 kb on either side to match the IMGT [24] definition, as all previously described *IGH::DUX4* rearrangements have breakpoints inside the *IGH* region [1]. Pelops uses two complementary *DUX4* region definitions while the previous implementation used only one to call all *DUX4* rearrangements. Both the core and the extended *DUX4* regions are different to the one previously used, which covered the *DUX4* repeat arrays with approximately 70 kb and 50 kb margins on chromosomes 4 and 10, respectively.

### Calling *DUX4*-rearrangements with non-*IGH* regions

Pelops’ method for calling *DUX4* fusions with non-*IGH* genes involves three main steps. First, candidate regions in the genome that have a potential translocation with *DUX4* are identified. Secondly, a slightly modified version of the algorithm developed for *IGH::DUX4* fusions is run for each of the candidate regions. Finally, candidate regions with low evidence based on the results of the previous step are filtered out.

#### Finding candidate regions

Candidate regions are found by looking for reads flagged as improperly paired in the core *DUX4* region. Only rearrangements with the core *DUX4* region are considered to increase specificity of this caller and to reduce noise from repetitive genomic elements located in the extended *DUX4* region. Mapping location of their mates are counted in 1 kb genomic region bins. Bins with more than two improperly paired reads are retained, and adjacent genomic bins are merged. All genomic bins overlapping with the extended *DUX4* and *IGH* regions are removed. This forms a provisional set of candidate genomic regions which may be rearranged with *DUX4*.

#### Finding evidence for rearrangements in each candidate region

For each candidate region, spanning reads are found in the same manner as described for *IGH::DUX4* fusions, except for one difference: spanning reads are only counted if the aligned segment in the candidate region has a mapping quality ≥ 10 (default threshold that can be changed by the user). This is to reduce the number of false positive rearrangements with candidate regions that contain repetitive sequences.

#### Filtering candidate regions

For the final output, blacklisted regions that were repeatedly called in normal samples, and regions with SRPB < 20 (default threshold that can be changed by the user) are removed. The blacklist used for filtering candidate regions contains all candidate regions that are called at least twice with SRPB ≥ 10 amongst the normal samples of the paediatric ALL cohort. As the blacklist is highly dependent on the read aligner used, separate blacklists were generated for DRAGEN and bwa analyses. The regions included in each blacklist are listed in Supplementary Table 5.

#### Validation of optimal parameters

The non-*IGH DUX4*-r caller is dependent on several parameters: the minimum mapping quality of reads in a candidate region, the minimum SRPB threshold for filtering candidate regions, and the minimum SRPB threshold used for generating the blacklist. To find the optimal settings, a random permutation cross validation was performed 10 times. The normal samples were randomly divided into a training (80%) and an evaluation (20%) set (the test set consists of the tumour samples). All candidate regions that were called at least twice in the training set, passing a given blacklist SRPB threshold, were added to the blacklist. Then, in the evaluation set, candidate regions above a given filtering SRPB threshold were compared against this blacklist. The effect of changing the minimum mapping quality threshold to 1 was also investigated in this way. This validation was done for both bwa and DRAGEN alignments. The final parameter values were chosen such that there were no additional calls in the evaluation set for any of the validation rounds, while yielding a minimal blacklist to minimise the risk of missing true positives in the test set.

### *De novo* assembly of breakpoints

As input for *de novo* assembly, spanning reads from DRAGEN-aligned BAM files were exported to a SAM file using Pelops. *De novo* assemblies were produced with SPAdes version 3.11.1 [26] with default parameters. Where this method failed, Velvet version 1.2.10 [27] was used for *de novo* assembly, with the following non-default parameters:

- K. The size of the k-mers used to create de Bruijn graphs in Velvet, was set to K = 21.
- Expected k-mer coverage. This was calculated from the average coverage over the entire genome reported by DRAGEN.
- Fragment length. The fragment length was calculated from the median insert length and read length reported by DRAGEN.

### Mapping of *de novo* assemblies to the reference genome

The assembled scaffolds were mapped to the GRCh38 reference genome using BLAT [28]. Per sample, up to 500 alignments with an alignment score ≥ 20 were evaluated and ranked by their alignment score. Next, BLAT results were manually filtered to remove duplicate alignments resulting from the repetitive nature of the *IGH* and *DUX4* genomic regions. In order of importance, the following criteria were used:

- The scaffold should be aligned as completely as possible to segments of the reference genome.
- The highest-scoring alignments should be used to cover each segment of the scaffold.
- Alignments should be consistent (e.g. for *DUX4,* all segments should be aligned to the same chromosome).

If multiple alignments were equivalent according to these criteria, the alignment most upstream on the reference genome was used.

In BLAT, each alignment can consist of multiple blocks. A custom Python script was used to extract these blocks and merge adjacent blocks if they differed only by short indels of length < 20 bp. Another custom Python script was used to annotate each aligned segment as *IGH*, core *DUX4,* extended *DUX4* (if not already covered by core *DUX4*), other, or unknown sequence (gaps in scaffold marked by Ns). Unannotated sequences > 25 bp were separately aligned with BLAT, with no minimum alignment score, to annotate short sequences that were previously missed.

From these alignments, a list of breakpoints was obtained. For our analyses, we only considered those breakpoints that have a continuous sequence in the *de novo* assembly, that is, without any unknown sequence connecting *IGH* and *DUX4* segments.

### Amplicon sequencing

Rearrangement junctions were confirmed by successful amplification of PCR products using primer pairs specific to each assembly. Primers were designed using *primer3* software [29–31] (Supplementary Table 6). PCR products were converted to sequencing libraries by combining 1 µl of PCR product with 19 µl of nuclease free water for input into the Nextera XT DNA Library Prep Kit (Illumina PN# 15032354, 15032355, 15052163). PCR amplicons were simultaneously fragmented and tagged then adapter sequences added by 12 cycles of PCR as per manufacturer’s protocol. Final library concentration was assessed on the Agilent 4200 TapeStation System using the High Sensitivity D1000 tape, and libraries were diluted to 2 nM with RSB and pooled. A 10 µl aliquot of pooled libraries was denatured with 10 µl freshly diluted 0.2 N NaOH (Sigma-Aldrich PN# 72068), through incubation for 5 minutes at room temperature. Following denaturation, the pooled library was then diluted to 20 pM by with addition of 980 µl of pre-chilled HT1 buffer (Illumina PN#15027041) and further diluted to a final loading concentration of 14 pM. The library was mixed with a 50% PhiX spike-in (Illumina PN# 15017397) and the final pool was sequenced on a MiSeq using a MiSeq reagent kit 150cy v3 (2x75bp reads) as per the manufacturer’s instructions (Illumina PN# 15043893, 15043894).

## Ethics approval and consent to participate

Not applicable.

## Consent for publication

Not applicable.

## Availability of data and materials

Pelops is publicly available as a Python package named ilmn-pelops. The results of this paper are based on Pelops version 0.8.0, which is available through the Python Package Index (https://pypi.org/project/ilmn-pelops/0.8.0/). The source code can also be found at https://github.com/Illumina/Pelops. The code has extensive unit test coverage (98.1% at commit 6f29f33e) and tests pass on Python 3.7, 3.9, 3.11. Pelops has minimal dependencies on third-party packages, with the exception of pysam [32]. The *IGH::DUX4* caller will also be implemented in Illumina DRAGEN 4.3.

## Authors’ contributions

PG and MM provided guidance for software development, evaluated Pelops on the paediatric patient cohort and wrote the manuscript. SB, KJC, PG, and PJC developed the Pelops software. JFP conceived the methods for calling *DUX4*-rearrangements. DJM designed PCR primers for amplicon sequencing of the rearrangement junctions, while CF, AI and HN performed and optimised PCR and sequencing. SLR, CJH, and AVM provided patient samples and orthogonal data. JB, MTR, SLR, CJH, and AVM revised the manuscript. SLR, DRB, CJH, AVM and MTR conceived the study. All authors read and approved the final manuscript.

## Acknowledgements

Primary childhood leukaemia samples used in this study were provided by the VIVO Biobank for Children and Young People with Cancer. We also thank all the members of the NCRI Childhood Cancer and Leukaemia Group (CCLG) Leukaemia Subgroup for access to material and data on clinical trial patients.

## Tables

See separate Excel sheet for Supplementary Tables.

