## Supplementary Figures for "A dedicated caller for *DUX4* rearrangements from whole-genome sequencing data"

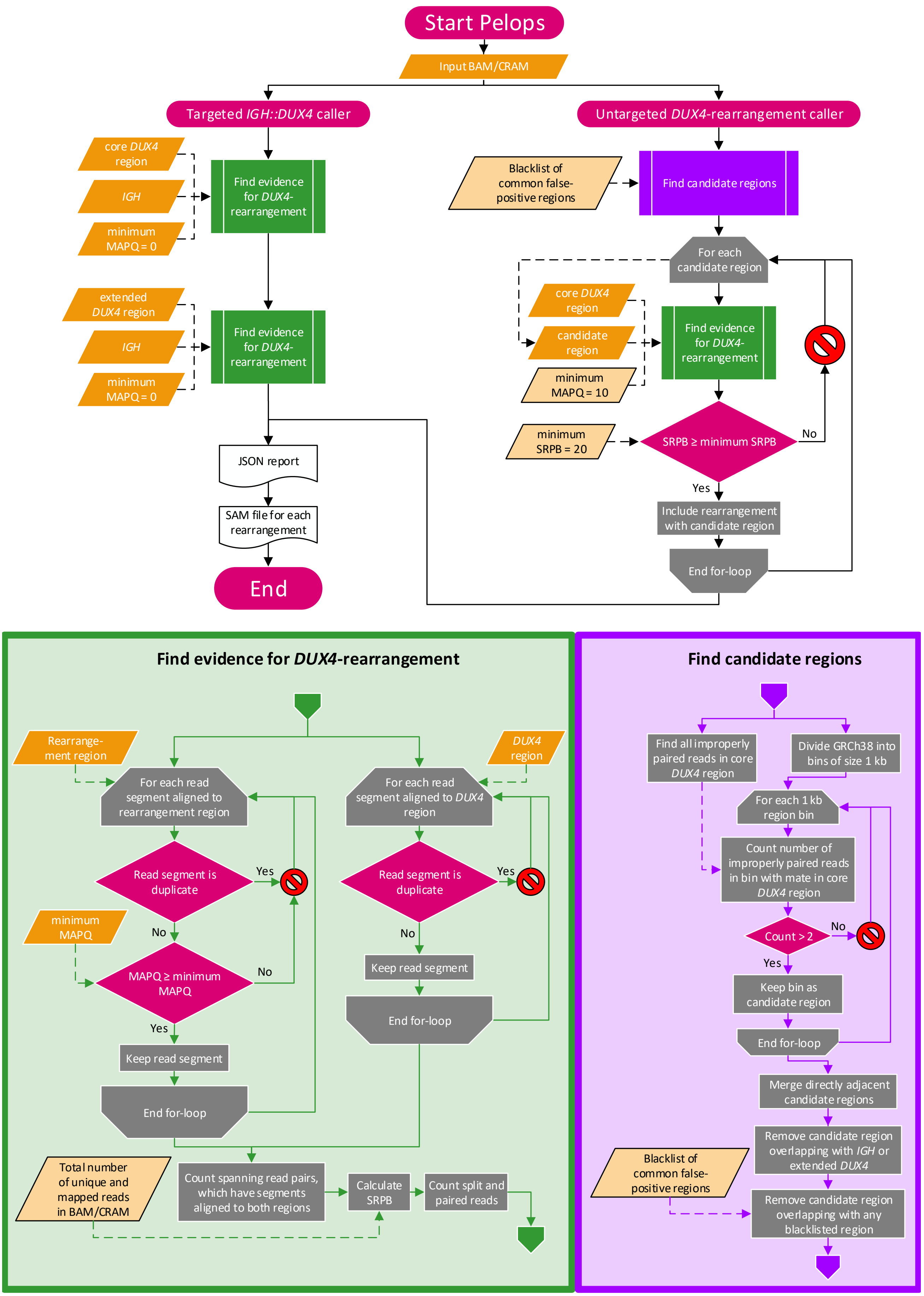

*Supplementary Figure S1*. Flowchart overview of Pelops. This overview illustrates the algorithm used by Pelops for calling *DUX4*-rearrangements, and the inputs, outputs and parameters of the pipeline. All parameters in the light orange parallelograms with a black frame can be changed by the user of Pelops through its command line interface.

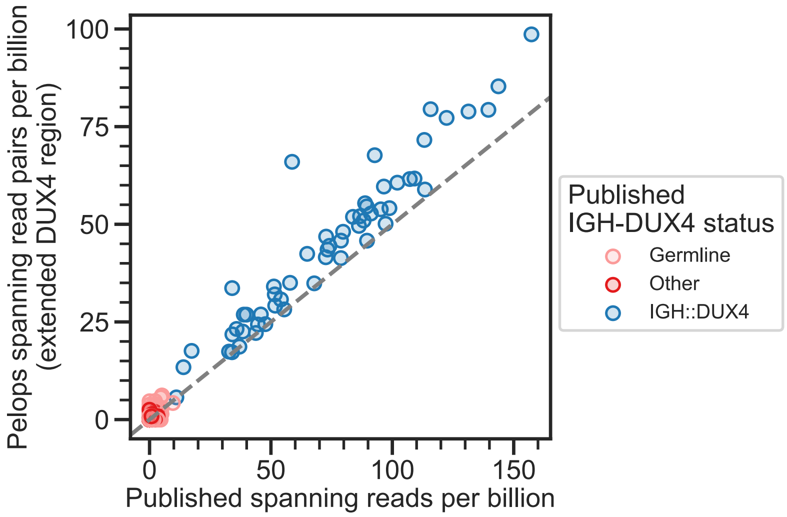


*Supplementary Figure S2*. Comparison of Pelops to previously published implementation (Ryan et al., 2023) using 210 tumour and 208 matched germline samples for the paediatric ALL cohort on Isaac-aligned BAM files. Colours indicate *DUX4*-r status of the sample, as previously published. The line indicates the expected difference based on counting spanning read pairs in Pelops versus the previously published method, which counted spanning reads individually.

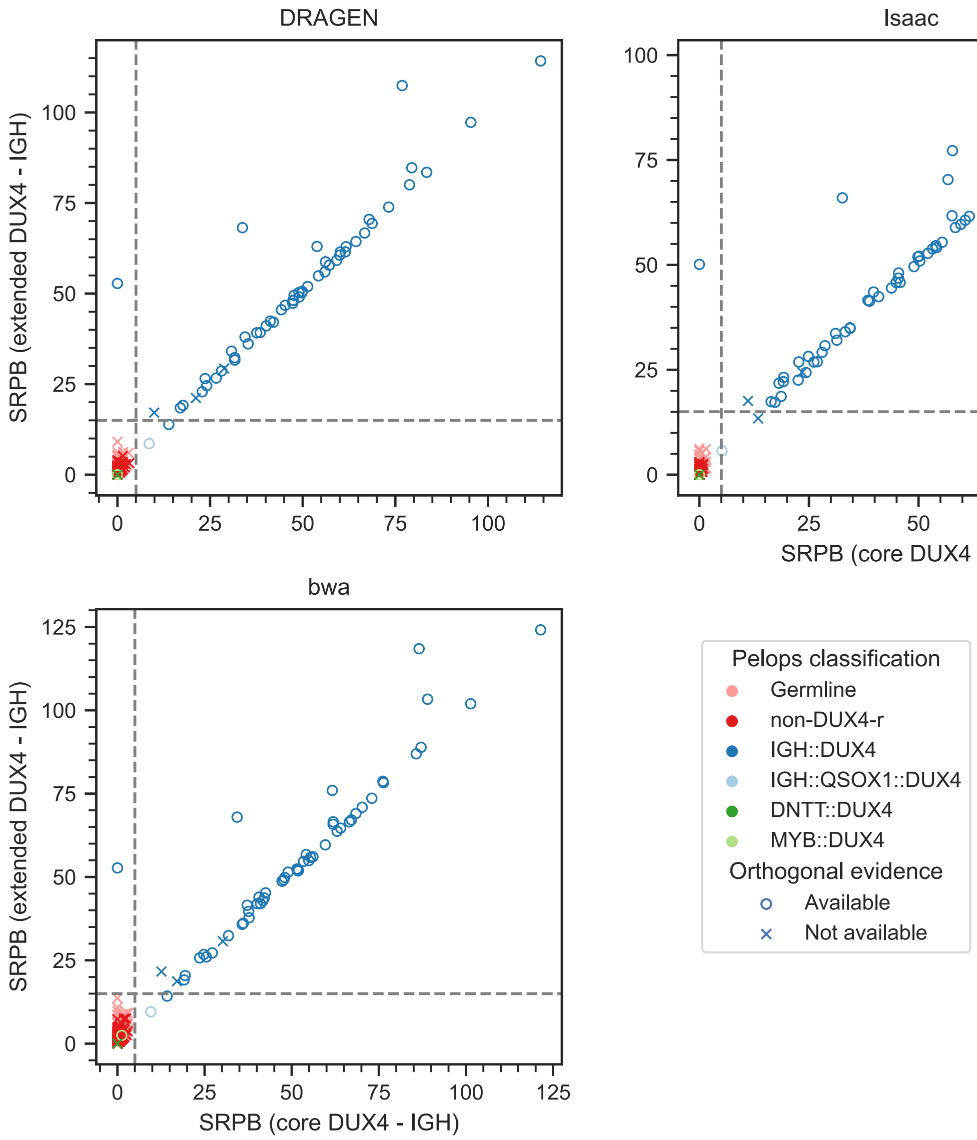


*Supplementary Figure S3*. Comparison of Pelops' *IGH::DUX4* caller results for three different aligners. Spanning read pairs per billion (SRPB) for *IGH::DUX4* were calculated by Pelops for all 210 tumour and 208 germline samples of the paediatric ALL validation cohort, using read alignments created by DRAGEN, bwa, and Isaac. The x-axis shows the SRPB distribution calculated based on the core *DUX4* region definition, while on the y-axis calculations are based on the extended *DUX4* region definition. Colours indicate *DUX4* fusion status of the samples predicted by Pelops. The dashed horizontal and vertical lines indicate SRPB thresholds of 15 and 5, for extended and core *DUX4* regions, respectively. The marker indicates whether orthogonal evidence for *DUX4*-r is available, based on RNA sequencing, the presence of *ERG* deletions, or amplicon sequencing.

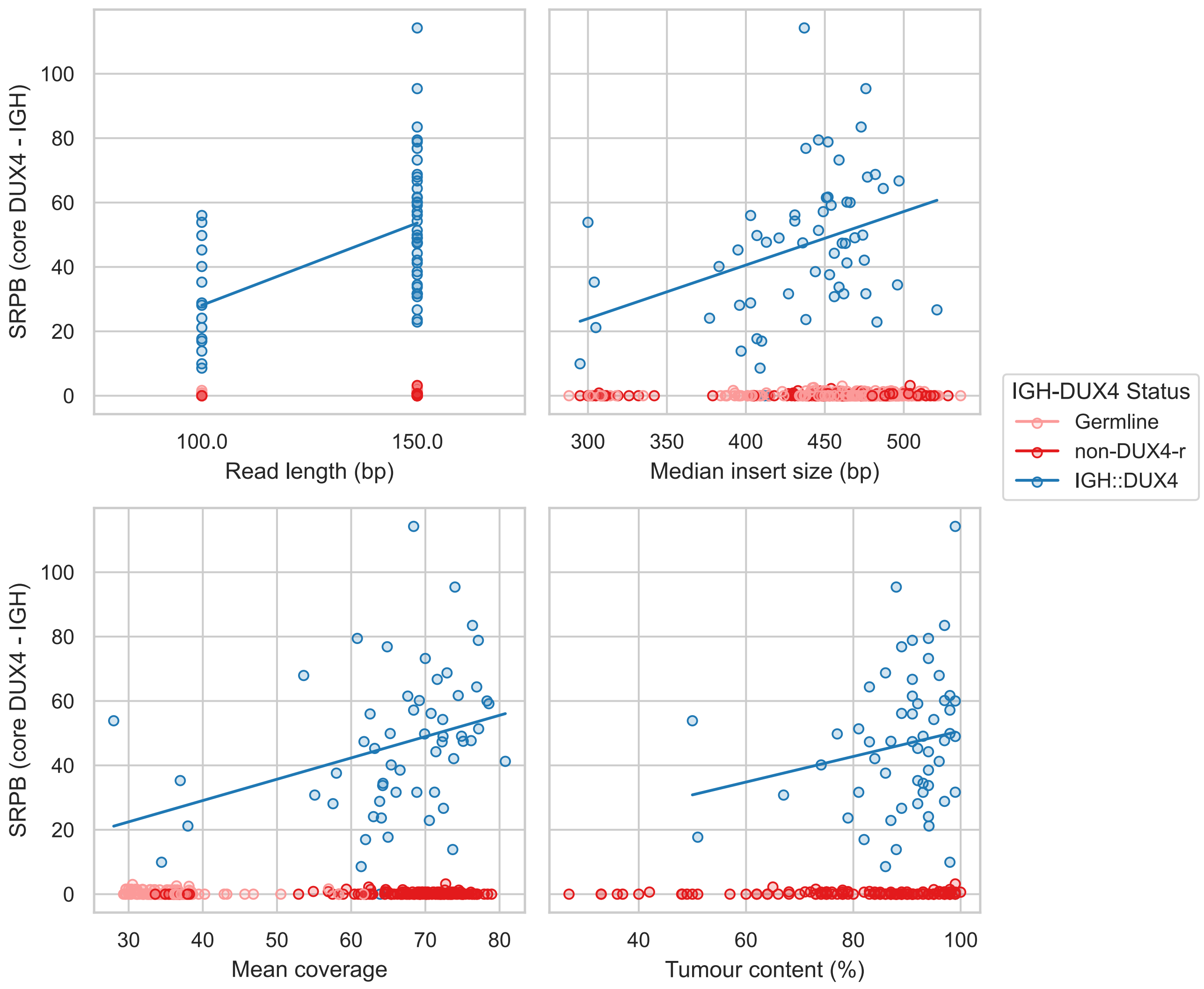


*Supplementary Figure S4.* Spanning read pairs per billion (SRPB) are plotted against the following sequencing parameters: read length, median insert size, mean coverage over genome and tumour content estimated by DRAGEN.

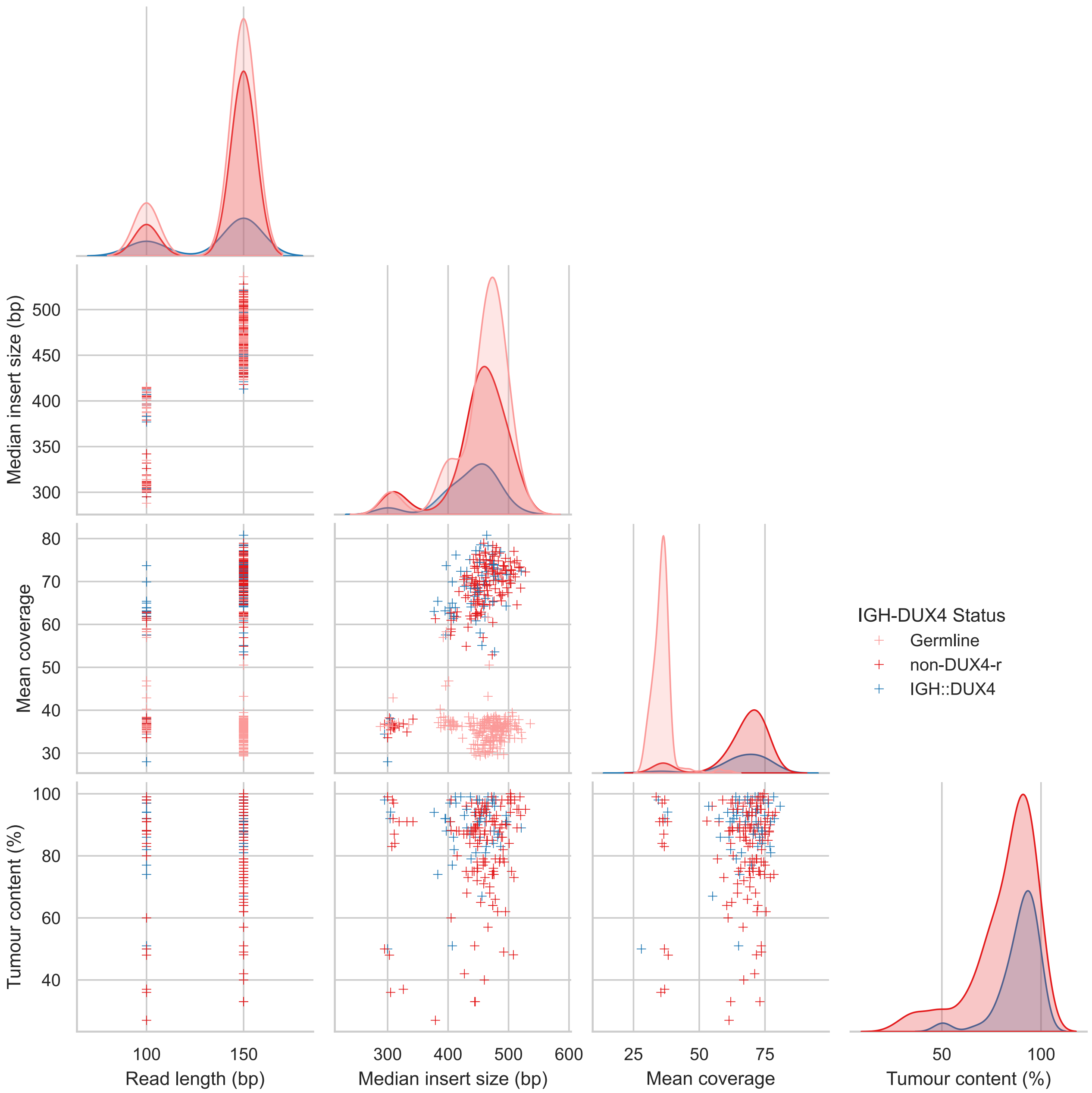


*Supplementary Figure S5.* Pair plot showing relationships between read length, median insert size, average alignment coverage over genome, and estimated tumour content in cohort samples. Off-diagonal scatterplots show relationships between two variables, while the on-diagonal plots visualise distributions of a single variable.

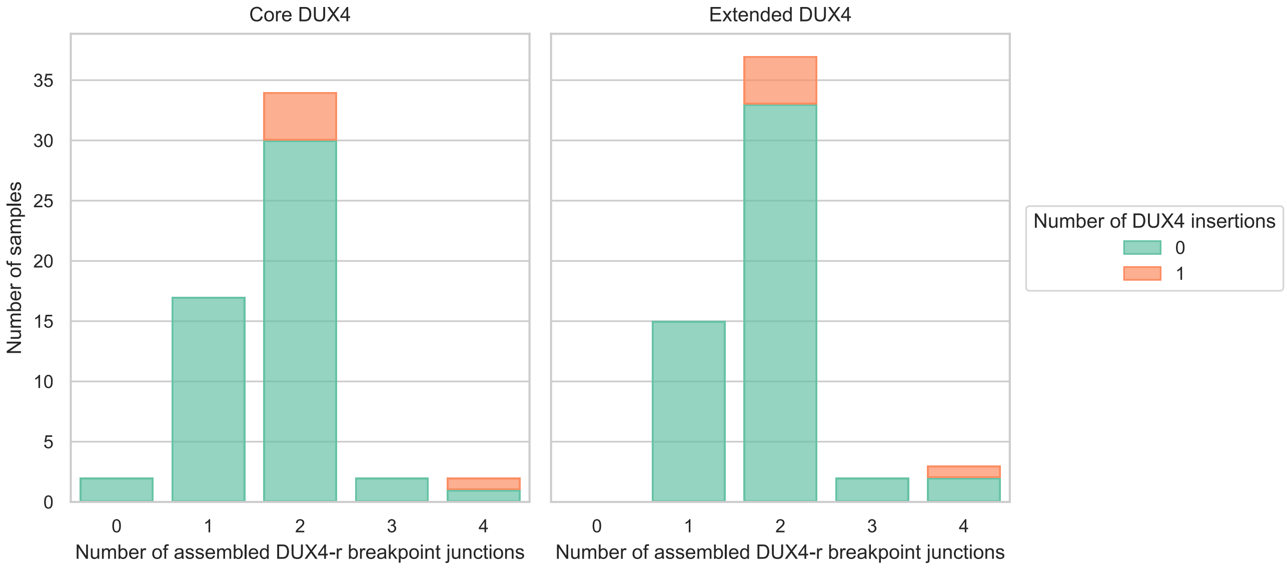


*Supplementary Figure S6*. Number of *IGH::DUX4* breakpoint junctions per sample in the *de novo* assemblies. Only junctions which have a continuously assembled sequence are counted. The left plot shows the number of junctions in the core *DUX4* region, the right plot the numbers in the extended *DUX4* region. Samples for which a full insertion of *DUX4* is observed are highlighted. Note that each insertion is counted as two separate breakpoint junctions.

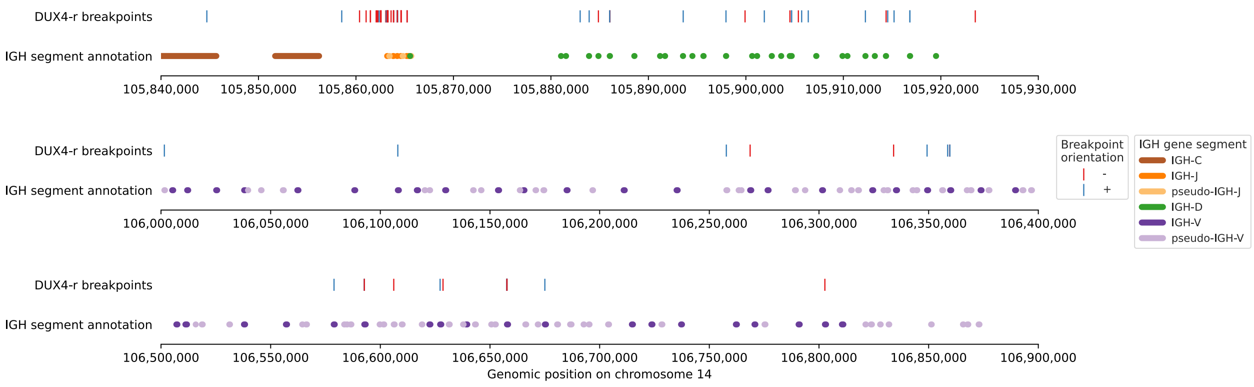


*Supplementary Figure S7*. Genomic location of *DUX4*-r breakpoints in the *IGH* locus for the paediatric patient cohort. Breakpoint orientation + means that the *DUX4* segment is downstream of the breakpoint, while – means that the *DUX4* segment is upstream of the breakpoint. Genomic positions are based on the GRCh38 reference. Segments of the *IGH* locus not shown here do not contain any *DUX4*-r breakpoint.
